# Stress granule component TIA-1 is a negative regulator of the non-canonical NLRP3 inflammasome

**DOI:** 10.1101/2025.07.12.634801

**Authors:** Prem Prasad Lamichhane, Aditi, Blake H. Neil, Paul B. Kilgore, Alfredo G. Torres, Ashok K. Chopra, Parimal Samir

## Abstract

Inflammasomes are cytosolic signaling hubs assembled upon pathogen- or damage associated molecular patterns (PAMP and DAMP) sensing by innate immune pattern recognition receptors (PRR). Lipopolysaccharide (LPS) present on the cell wall of gram-negative bacteria is a PAMP that activates caspase 11 (CASP11) dependent nucleotide-binding oligomerization domain-like receptor pyrin domain-containing 3 (NLRP3) inflammasome (known as non-canonical NLRP3 inflammasome) leading to pyroptosis. Several host factors are shown to promote non-canonical NLRP3 inflammasome activation by making LPS readily available for recognition by CASP11. Here, we report T-cell intracellular antigen-1 (TIA1), an RNA binding protein as a negative regulator of non-canonical NLRP3 inflammasome. Using bone marrow-derived macrophages (BMDMs), we demonstrated that the loss of TIA1 led to an increase in caspase-1 (CASP1) activity in response to cytosolic LPS. A previous study had demonstrated that mice lacking *Tia1* are more susceptible to LPS mediated endotoxic shock. Our results provide a potential explanation for this observation by showing loss of TIA1 increases non-canonical NLRP3 inflammasome activation resulting in increased inflammation and pathogenesis during LPS mediated endotoxic shock. Further, TIA1 mediated inhibition of non-canonical NLRP3 inflammasome is independent of TIA1’s regulatory role in gene transcription as well as its role in stress granule assembly. TIA1 is also dispensable for activation of the canonical NLRP3 inflammasome as well as AIM2 and NLRC4 inflammasomes. While, the exact mechanism by which TIA1 inhibits non-canonical inflammasome activation remains to be elucidated, our finding that TIA1 is a negative regulator indicates the presence of undiscovered regulatory mechanisms. Future studies will focus on unraveling these mechanisms for developing anti-inflammatory drugs that exploit non-canonical inflammasome activity modulation.

## Introduction

Inflammasomes are cytosolic, multiprotein signaling complexes that usually consist of a pattern recognition receptor (PRR) sensor, an adapter (apoptosis-associated speck-like protein containing a CARD (ASC), and an effector CASP1) (1,2). Different sensors, such as nucleotide-binding oligomerization domain-like receptor (NLR) pyrin domain containing 3 (NLRP3), NLRP1, NLR family CARD containing protein 4 (NLRC4), absent in melanoma 2 (AIM2), and PYRIN can sense diverse PAMPs and DAMPs (2). The extensively studied canonical NLRP3 inflammasomes are activated by a wide range of stimuli. In contrast, activation of the noncanonical NLRP3 inflammasome initiates with CASP11 (CASP4/5 in humans) that directly senses intracellular lipopolysaccharide (LPS) (3). Oligomerized CASP11 becomes autoproteolytically activated and cleaves Gasdermin D (GSDMD). The amino terminal fragment of GSDMD translocates to the plasma membrane and form pores through which K^+^ ion efflux happens, resulting in activation of the NLRP3 inflammasome (4). Wang *et al.,* provided the proof of concept that loss of CASP11 can protect from lethal endotoxic shock mediated by intraperitoneal injection of LPS. Further, they suggested CASP1 is downstream of CASP11 (5). The mechanism behind the critical function of CASP11 in propagating the LPS-triggered inflammatory response remained unexplored for more than a decade. Seminal studies by Kayagaki *et al.* (3), Shi et al (6), and Hagar *et al.* (7), demonstrated that cytosolic LPS is the ligand for CASP11, resulting in inflammasome activation and pyroptosis.

Various host factors tightly regulate non-canonical NLRP3 inflammasome activation at several steps. High mobility group box 1 binds to LPS facilitating its uptake by macrophages and subsequent release into the cytosol to activate CASP11 (8). Guanylate-binding proteins (GBPs) and immunity-related GTPase B10 (IRGB10) together, target the cell membranes of intracellular bacteria (9), or phagocytosed outer membrane vesicles derived from extracellular bacteria (10), causing the release of LPS into the cytosol leading to CASP11 activation. Additionally, toll-like receptor 4 and its adapter TIR-domain-containing adapter-inducing interferon-β (TRIF) are essential for promoting CASP11 expression and activation in response to enteropathogens such as *Citrobacter rodentium* and *Escherichia coli* (11,12). These studies have revealed host factors that either promote the release of LPS for sensing by CASP11 or enhance CASP11 expression.

Host factors that suppress activation of the non-canonical NLRP3 inflammasome remain elusive and understudied. One potential negative regulator of the non-canonical NLRP3 inflammasome is T-cell intracellular antigen-1 (TIA1), which is a RNA-binding protein (RBP), and plays a key role in post-transcriptional gene expression regulation (13–17). TIA1 was reported to dampen translation of tumor necrosis factor (TNF) mRNA to protect against hyper-inflammation (13,17). In support of this thesis, heightened production of TNF has been attributed to the hypersensitivity of *Tia1*^-/-^ mice to LPS induced endotoxic shock (17). However, in the same seminal paper, authors anticipated that an alternative explanation for their results was possible and it mentioned in the manuscript (17). The non-canonical NLRP3 inflammasome was not discovered at the time of publication. Although it was known that *Casp11*^-/-^ mice were resistant to LPS mediated endotoxic shock (5), only later studies showed that the non-canonical NLRP3 inflammasome was assembled by activation of CASP11 (CASP4/5 in humans) upon intracellular LPS sensing, thus providing a mechanistic explanation for observed results in animal experiments. (4,6,7). Therefore, in this study we investigated the effect of TIA1 loss on non-canonical NLRP3 inflammasome activation. We found that TIA1 plays a critical role in suppressing non-canonical NLRP3 inflammasome activation, highlighting the importance of TIA1 in host defense during sepsis.

## Materials and Methods

### Mice

B6.129S2(C)-*Tia1tm1Andp*/J mouse (17) (Jackson Laboratory, 009248) and BL6/J wild type mice were acquired from the Jackson Laboratories. All mice were bred at the University of Texas Medical Branch (UTMB) at Galveston. Animal studies were carried out in accordance with protocols approved by the UTMB Institutional Animal Care and Use Committee.

### Bone marrow-derived macrophage culture and stimulation

Primary BMDMs were cultured for 6 days in DMEM (Sigma-Aldrich, D5671) supplemented with 10% fetal bovine serum (Biowest, S1620), 30% L929-conditioned medium (the base medium is IMDM, Sigma-Aldrich, I3390), 1% non-essential amino acids (Sigma-Aldrich, M7145) and 1% penicillin and streptomycin (Sigma-Aldrich, P4333). The culture was supplemented with BMDM media on day 3 and 5. BMDMs were seeded at a density of one million cells per well in a 12-well plate and incubated overnight. BMDMs were washed with warm PBS and incubated with warm DMEM supplemented with 10% FBS for 30 min before use.

For NLRP3 inflammasome activation, BMDMs were primed with 100 ng/mL of Ultrapure LPS (*Salmonella* Minnesota *R595*; tlrl-smlps; InvivoGen) for 4 h, 50 µg/mL of low molecular weight poly(I:C) (InvivoGen, tlrl-picw) for 6 h, 1 µg/mL of R848 (resiquimod) (tlrl-r848-5; InvivoGen) for 6 h, or 1 µg/mL of Pam3CSK4 (tlrl-pms; InvivoGen) for 6 h followed by addition of adenosine triphosphate (ATP, final concentration of 5 mM) (Cayman Chemicals Inc., 14498) for 30 to 45 min.

For AIM2 activation, BMDMs in Opti-MEM (Thermo Fisher Scientific, 31985-070) were transfected for 2 h with 1 µg/well of poly(dA:dT) (InvivoGen, tlrl-patn) mixed with Xfect polymer (Takara Bio, ST0154) as per manufacturer’s instructions.

For non-canonical inflammasome activation, BMDMs in Opti-MEM were primed with 1 µg/mL of Pam3CSK4 (tlrl-pms; InvivoGen) for 2 h or left unprimed. The cells were transfected for 3 h with 2 µg/well of Ultrapure LPS (*S.* Minnesota *R595*; tlrl-smlps; InvivoGen) mixed with Xfect polymer (Takara Bio, ST0154) as per manufacturer’s instructions.

For bacterial infection mediated inflammasome activation, BMDMs were infected with *S. enterica* serovar Typhimurium 14028s (multiplicity of infection (MOI) 1) in antibiotic free complete DMEM. Cells lysates were harvested when approximately 50% of WT cells showed pyroptotic morphology. *Aeromonas dhakensis* (MOI 20) (18)*, Citrobacter rodentium* DBS100 (19) (MOI 10), or *Escherichia coli* HS (20) (MOI 10) in antibiotic free complete DMEM, centrifuged (RCF = 500 g) for 5 minutes, and after two hours cells were washed with warm pbs and complete DMEM containing 50 µg/mL gentamicin was added. *A. dhakensis* infected cell lysates were harvest when approximately 50% of WT cells showed pyroptotic morphology. *C. rodentium* and *E. coli* infected lysates were harvested after overnight incubation.

### Western blot analysis

For caspase-1 immunoblotting, BMDMs and the culture supernatants were combined in caspase lysis buffer (containing protease inhibitors, 5% NP-40 and 25 mM dithiothreitol) and sample-loading buffer (containing SDS and 2-mercaptoethanol). For signaling immunoblot, BMDMs were lysed in 1X RIPA and sample loading buffer. Protein samples were boiled for 5 min at 95 ⁰C followed by separation on 8-12% polyacrylamide gels. The proteins were transferred onto PVDF membranes (EMD Millipore, IPVH00010) followed by blocking in 5% skim milk for 1 h at room temperature. The blocked membranes were incubated overnight at 4 ⁰C with primary antibodies. The primary antibodies were anti-CASP1 (AdipoGen, AG-20B-0042), anti-CASP11 (SantaCruz, sc-56038), anti-CASP11 (Novus Biologicals, NB120-10454), anti-GSDMD (Abcam, ab209845), anti-NLRP3 (Adipogen, AG-20B-0014-C100), anti-ASC (Adipogen, AG-25B-0006-C100), anti-cleaved-IL1b (CST, 63124), anti-GAPDH (Proteintech, 60004-1-Ig), and CoraLite Plus 750-conjugated anti-beta actin (Proteintech, CL750-66009). The membranes were washed three times with 1X TBS containing 0.05% Tween-20 followed by incubation with secondary horseradish peroxidase-conjugated antibodies ([315-035-047; anti-mouse]; [anti-rabbit,111-035-047]; [anti-rat, 112-035-006]; Jackson ImmunoResearch Laboratories) for 1 h. Protein bands were visualized using Luminata Forte Western HRP Substrate (Millipore, WBLUF0500) and the images were acquired on ChemiDoc (BioRad) and analyzed using ImageJ Fiji v1.54.

### Confocal Microscopy Imaging and analysis

BMDMs were seeded at a density of eighty thousand cells per well in 18-well chambered cover glass (C18-1.5H, Cellvis, Mountain View, CA). After overnight incubation, the cells were washed with warm PBS and incubated with warm DMEM supplemented with 10% FBS for 30 min prior to stimulation. The BMDMs in Opti-MEM were primed with 1 µg/mL of Pam3CSK4 (tlrl-pms; InvivoGen) for 2 h or left unprimed. The cells were transfected with 200 ng/well of Ultrapure LPS (*S.* Minnesota *R595*; tlrl-smlps; InvivoGen) mixed with Xfect polymer (Takara Bio, ST0154) as per manufacturer’s instructions for the indicated time. Additionally, the unprimed cells were stimulated with 10 µg/mL of Ultrapure LPS (*Salmonella* Minnesota *R595*; tlrl-smlps; InvivoGen).

The cells were processed as described previously (21). Briefly, the stimulated cells were fixed in 4% paraformaldehyde (sc-281692, Santa Cruz Biotechnology Inc.) at room temperature for 15 min and washed with PBS followed by blocking in 2% BSA in PBS. BMDMs were stained with anti-ASC (1:250, Adipogen, AG-25B-0006-C100) and anti-G3BP1 (1:1000; Proteintech; 66486-1-Ig) overnight at 4 °C. The cells were washed 3 times in PBS and incubated with the Alexa Fluor 555-conjugated anti-rabbit IgG (1:1000; Life Technologies; A32794), Alexa Fluor 488–conjugated anti-mouse IgG (1:1000; Life Technologies; A32766) secondary antibody, and DAPI (5 μg/mL, Cayman Chemicals Inc.). Confocal images were acquired using an ImageXpress® Micro Confocal (Molecular Devices, San Jose, CA, USA) confocal microscope. Images were analyzed using ImageJ Fiji v1.54. Stress granules and ASC were detected using ilastik (22) and CellProfiler (23) software using an integrated pipeline described in (24).

### Reverse transcription quantitative PCR analysis

Following the stimulations, cells were lysed in TRIZOL (Thermo Fisher Scientific, 15596026) at indicated time-points followed by RNA extraction according to the manufacturer’s protocol. 250 ng of RNA was reverse transcribed using High-Capacity cDNA Reverse Transcriptase kit (Applied Biosystem, 4368813). Quantitative PCR was performed on the BioRad CFX Connect^TM^ real-time PCR instrument using 2× SYBR Green (Applied Biosystems, 4368706). The primers used are as follows: *Gapdh*: 5′-CGT CCC GTA GAC AAA ATG GT-3′, 5′-TTG ATG GCA ACA ATC TCC AC-3′; *Il1b*: 5′-GAT CCA CAC TCT CCA GCT GCA-3′, 5′-CAA CCA ACA AGT GAT ATT CTC CAT G-3′; *Tia1*: 5′-TGG GCG GAA GAT AAT GGG TAA G-3′, 5′-GCT CGT ATC CTT CTT TTG ACT GC-3′; *Casp11*: 5′-ACA AAC ACC CTG ACA AAC CAC-3′, 5′-CAC TGC GTT CAG CAT TGT TAA A-3′; *Nlrp3*: 5′-ATT ACC CGC CCG AGA AAG G-3′, 5′-TCG CAG CAA AGA TCC ACA CAG-3′; *Casp1*: 5′-ACA AGG CAC GGG ACC TAT G-3′, 5′-TCC CAG TCA GTC CTG GAA ATG-3′; *Asc*: 5′-CTT GTC AGG GGA TGA ACT CAA AA-3′, 5′-GCC ATA CGA CTC CAG ATA GTA GC-3′; *Gbp2*: 5′-CTG CAC TAT GTG ACG GAG CTA-3′, 5′-GAG TCC ACA CAA AGG TTG GAA A-3′; *Gbp5*: 5′-CAG ACC TAT TTG AAC GCC AAA GA-3′, 5′-TGC CTT GAT TCT ATC AGC CTC T-3′; *Ifnb*: 5′-GCC TTT GCC ATC CAA GAG ATG C-3′, 5′-ACA CTG TCT GCT GGT GGA GTT C-3′; *Ninj1*: 5′-GAG TCG GGC ACT GAG GAG TAT-3′, 5′-CGC TCT TCT TGT TGG CAT AAT GG-3′; *Tnf*: 5′-CCC TCA CAC TCA GAT CAT CTT CT -3′, 5′-GCT ACG ACG TGG GCT ACA G-3′. The expression was normalized to the expression of *Gapdh*.

### Statistical analysis

GraphPad Prism v9 software was used for statistical analysis. Statistical significance of the data was determined by the paired *t*-test or two-way ANOVA method. Data are presented as mean ± SEM. *p* values less than 0.05 were considered statistically significant where ns is *p* > 0.05, ∗ *p* < 0.05, ∗∗ *p* < 0.01, ∗∗∗ *p* < 0.001 and ∗∗∗∗ *p* < 0.0001.

## Results

### TIA1 dampens non-canonical NLRP3 inflammasome activation

TIA1 loss has been shown to increase lethality in LPS mediated endotoxic shock in mice (17), highlighting its protective role during sepsis. To investigate whether the protective role of TIA1 could be mediated through regulation of non-canonical NLRP3 inflammasome during LPS mediated endotoxic shock, we transfected LPS into Pam3CSK4 primed or unprimed bone marrow-derived macrophages (BMDMs) isolated from WT and *Tia1*^-/-^ mice and measured CASP1 activation using western blot analysis. A technical challenge for detecting inflammasome activation increase is that if cells are incubated for a sufficient duration, all cells would undergo pyroptosis, which would mask increased kinetics of inflammasome activation. Therefore, we visually monitored cell death after LPS transfection and harvested the lysates when half of the WT cells in Pam3CSK4 primed condition were dead. With this approach, we observed increased abundance of the p20 fragment of CASP1 indicating stronger activation of non-canonical NLRP3 inflammasome in *Tia1*^-/-^ compared with WT BMDMs (**Fig. 1A-D**). Infection with enteropathogens like *C, rodentium* and *E. coli* also trigger non-canonical NLRP3 inflammasome activation (11,12).

**Fig 1:**
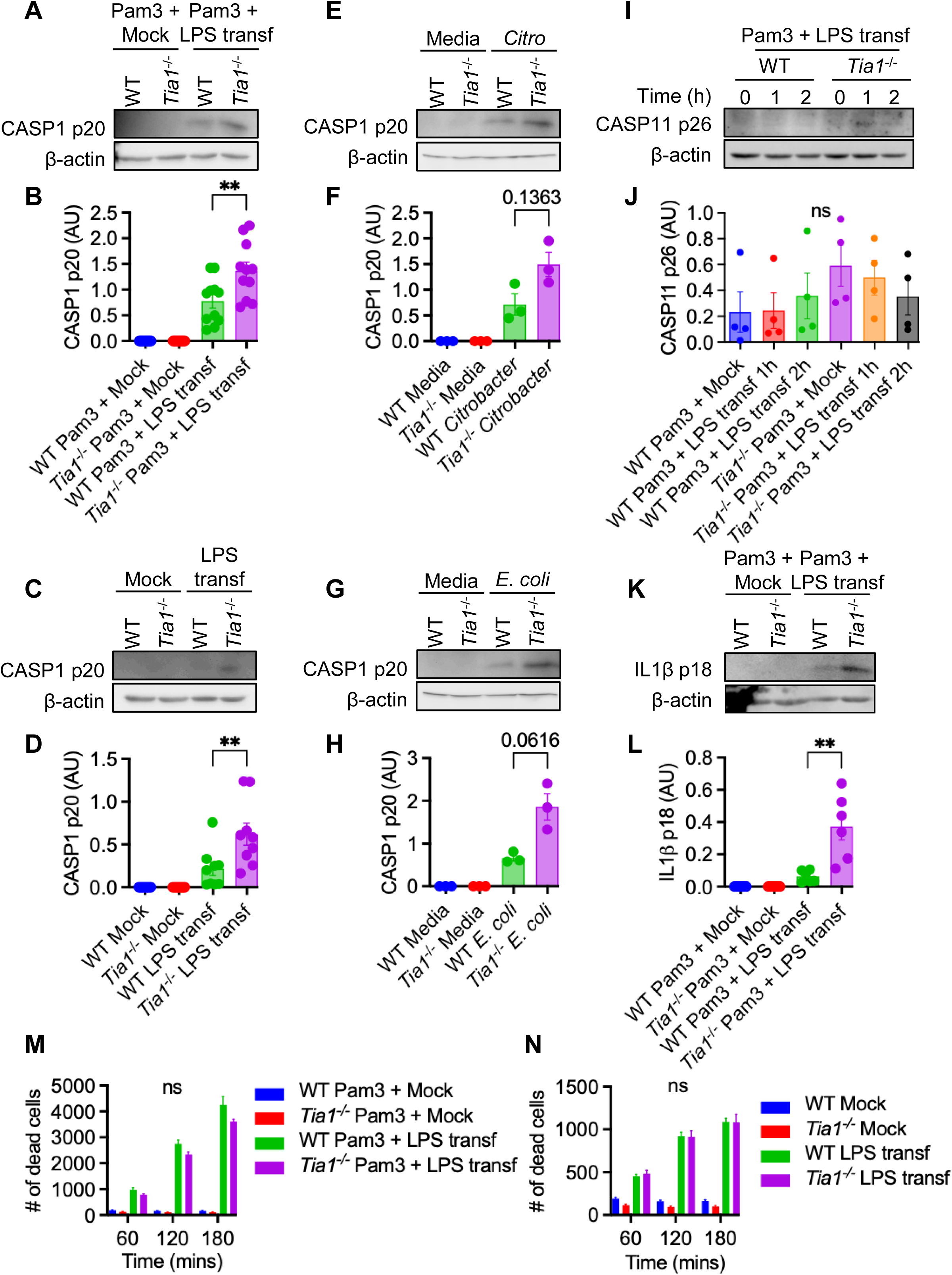
TIA1 dampens non-canonical NLRP3 inflammasome activation. **(A)** Immunoblot analysis of CASP1 with loading control b-actin in Pam3CSK4 primed BMDMs transfected with LPS for 2 hours (n = 11). **(B)** Densitometric quantification of CASP1 p20 bands in panel A. Values for CASP1 p20 are normalized to the loading control and expressed as arbitrary units (AU). **(C)** Immunoblot analysis of CASP1 with loading control b-actin in BMDMs transfected with LPS for 2 hours (n = 9). **(D)** Densitometric quantification of CASP1 p20 bands in panel C. Values for CASP1 p20 are normalized to the loading control and expressed as arbitrary units (AU). **(E)** Immunoblot analysis of CASP1 with loading control b-actin in BMDMs infected with *Citrobacter rodentium* at an MOI of 10 (n = 3). **(F)** Densitometric quantification of CASP1 p20 bands in panel E. Values for CASP1 p20 are normalized to the loading control and expressed as arbitrary units (AU). **(G)** Immunoblot analysis of CASP1 with loading control b-actin in BMDMs infected with *Escherichia coli* at an MOI of 10 (n = 3). **(H)** Densitometric quantification of CASP1 p20 bands in panel G. Values for CASP1 p20 are normalized to the loading control and expressed as arbitrary units (AU). **(I)** Immunoblot analysis of cleaved CASP11 with the loading control b-actin in BMDMs primed with Pam3CSK4 for 2 hours followed by LPS transfection (n = 4). **(J)** Densitometric quantification of CASP11 p26 bands in panel I. Values for CASP11 p26 are normalized to the loading control and expressed as arbitrary units (AU). **(K)** Immunoblot analysis of IL1b with loading control b-actin in Pam3CSK4 primed BMDMs transfected with LPS for 2 hours (n = 6). **(L)** Densitometric quantification of IL1b p20 bands in panel G. Values for CASP1 p20 are normalized to the loading control and expressed as arbitrary units (AU). **(M)** Sytox Green based measurement of RCD in Pam3CSK4 primed BMDMs transfected with LPS. **(N)** Sytox Green based measurement of RCD in unprimed BMDMs transfected with LPS Western blots are representative from at least three independent experiments. Data are shown as mean ± SEM (B, D, F, H, J, L, M and N). Statistical significance was evaluated using paired *t*-test (B, D, F, H, J, and L) or two-way ANOVA (M and N). ns, non-significant (*p* > 0.05), ∗*p* < 0.05 and ∗∗*p* < 0.01.

To investigate the role of *Tia1*^-/-^ in regulating non-canonical NLRP3 inflammasome activation during infection with enteric bacteria, we infected WT and *Tia1*^-/-^ BMDMs with *C*. *rodentium* and *E*. *coli* at a MOI of 10, and measured CASP1 activation using western blot analysis. We observed a trend towards increased CASP1 activation following infection with *C*. *rodentium* (**Fig. 1E-F**, 2.09-fold increase, *p*=0.1363) and *E*. *coli* (**Fig. 1G-H**, 2.79-fold increase, *p*=0.0616) suggesting *Tia1*^-/-^ negatively regulates non-canonical NLRP3 inflammasome activation following infection with the enteropathogens as well. During the non-canonical NLRP3 inflammasome signaling cascade, CASP1 activation is downstream of CASP11 activation (25). Although, we did not find a statistically significant increase in the abundance of the CASP11 p26 fragment, there is a trend towards abundance in *Tia1*^-/-^ compared to WT BMDMs (**Fig. 1I-J**). CASP1 cleaves pro-IL-1β into the biologically active p18 fragment. Supporting the role of TIA1 as an inhibitor of the non-canonical inflammasome, we observed an increase in the abundance of cleaved IL-1β p18 fragment in *Tia1*^-/-^ BMDMs compared to WT BMDMs (**Fig. 1K-L**) upon non-canonical NLRP3 inflammasome activation in Pam3CSK4 primed BMDMs. Interestingly, we did not observe any difference in cell death kinetics in WT and *Tia1*^-/-^ BMDMs (**Fig. 1M-N**) after LPS transfection. ASC, an adaptor protein and a component of NLRP3 inflammasome, oligomerizes to form a microscopically visible cytoplasmic supramolecular structure called ASC-specks. ASC specks amplify inflammasome activation most likely by providing a platform for more efficient processing of CASP1 into the active p20 fragment (26). Therefore, we performed immunofluorescence confocal microscopy to visualize and quantify ASC-speck formation in Pam3CSK4 primed or unprimed WT and *Tia1*^-/-^ BMDMs transfected with LPS. We observed significantly more ASC specks in *Tia1*^-/-^ BMDMs than WT BMDMs (**Fig. 2**). As expected, Pam3CSK4 priming increased the number of ASC positive BMDMs upon LPS transfection. Together, these results suggest that TIA1 dampens non-canonical NLRP3 inflammasome activation, ASC speck assembly, and IL-1β processing.

**Fig. 2:**
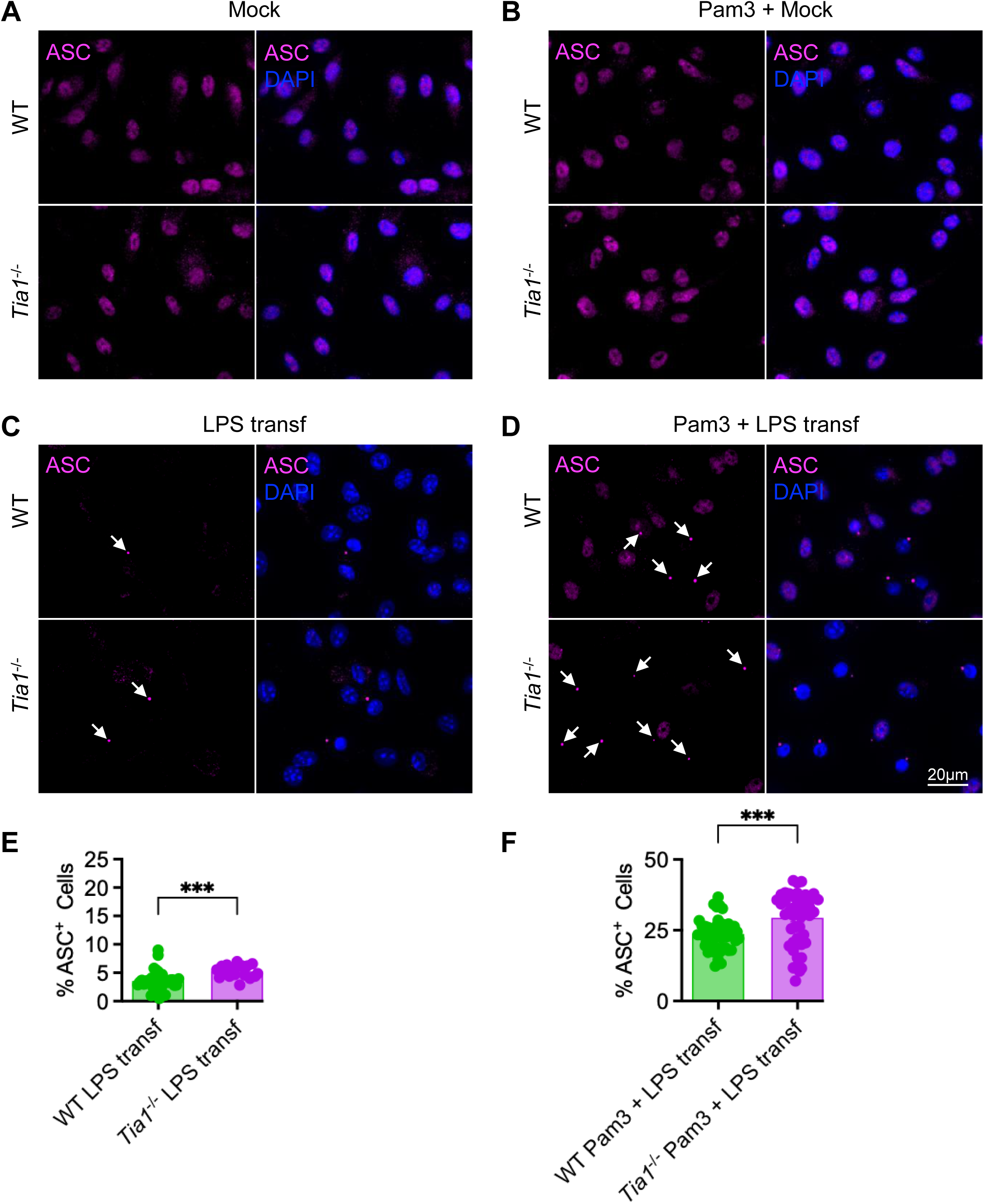
TIA1 inhibits ASC speck assembly downstream of non-canonical NLRP3 inflammasome activation. **(A-D)** Confocal microscopy of unprimed BMDMs (A and C) or BMDMs primed with Pam3CSK4 (B and D) for 2 hours followed by LPS transfection (C-D) for 120 min. (E-F) Quantification of ASC speck (E) in panel C and (E) in panel D. Blue: DAPI, Magenta: ASC. The data are representative of at least three independent experiments. Statistical significance was evaluated using unpaired *t*-test. ∗∗∗*p* < 0.001.

### TIA1 is dispensable for canonical NLRP3 inflammasome activation

Activation of CASP11 and subsequent cell death, unlike canonical activation of CASP1 and pyroptosis, does not require NLRP3 and ASC (25). Therefore, we next determined whether the observed TIA1-mediated decrease in CASP1 activation is specific to non-canonical triggers or if it is a general phenomenon happening with canonical triggers as well. Comparable levels of CASP1 activation were observed between WT and *Tia1*^-/-^ BMDMs treated with LPS plus ATP (**Fig. 3A-B**), Pam3CSK4 plus ATP (**Fig. 3C-D**), polyinosinic:polycytidylic acid (poly(I:C)) plus ATP (**Fig. 3E-F**) and R848 plus ATP (**Fig. 3G-H**). Because some gram-negative bacteria, including *Aeromonas dhakensis* (27) can induce robust canonical NLRP3 inflammasome activation, we investigated the effect of TIA1 loss on CASP1 activation upon infection with *A. dhakensis*. We found comparable activation of CASP1 in WT and *Tia1*^-/-^ BMDMs infected with *A. dhakensis* (**Fig. 3I-J**). Overall, these observations suggested that TIA1 is dispensable for canonical NLRP3 inflammasome activation.

**Fig 3:**
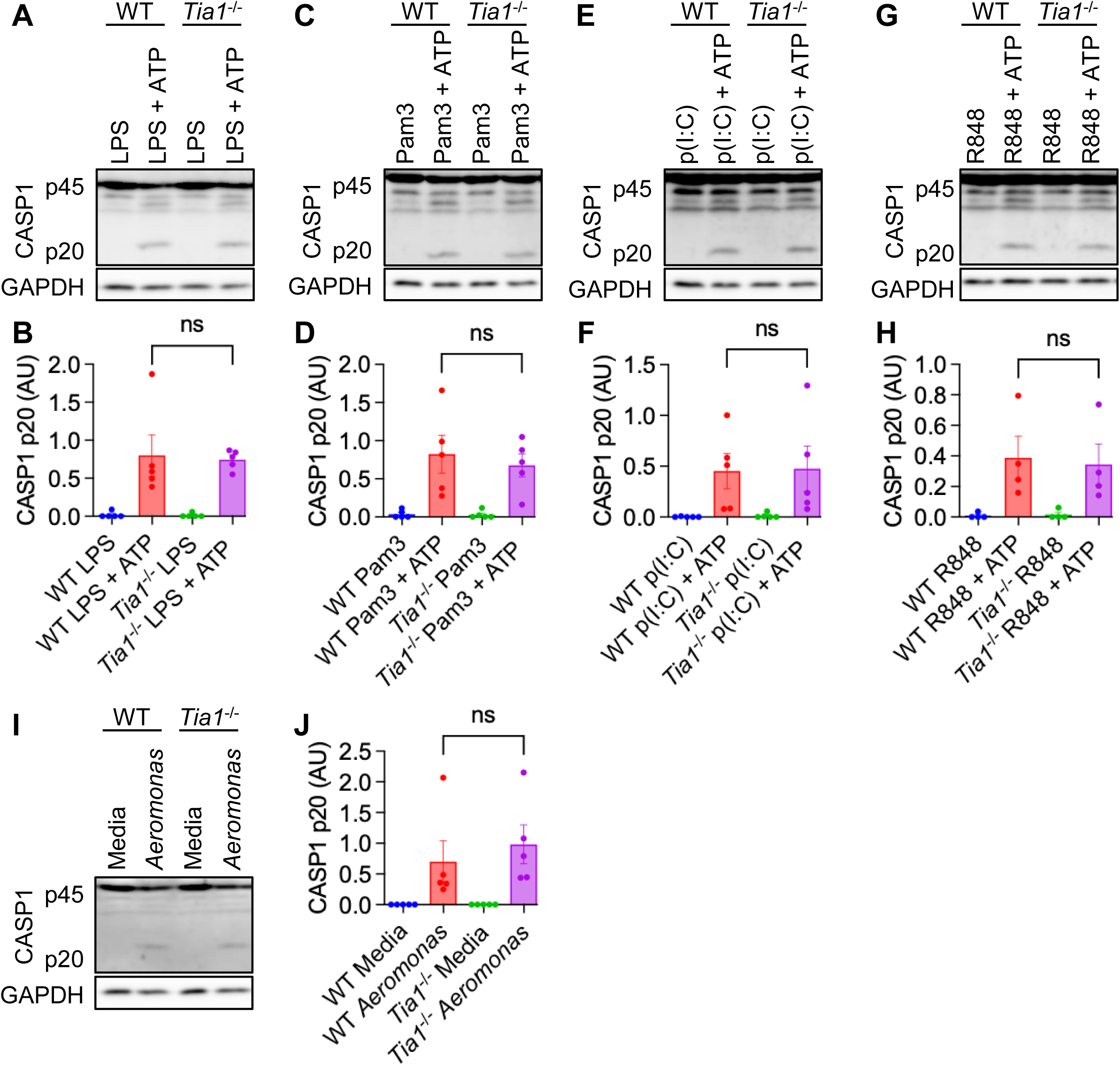
TIA1 is dispensable for canonical NLRP3 inflammasome activation. **(A)** Immunoblot analysis of CASP1 with loading control GAPDH in BMDMs primed with LPS for 4 hours followed by ATP treatment for 45 minutes (n = 5). **(B)** Densitometric quantification of CASP1 p20 bands in panel A. Values for CASP1 p20 are normalized to the loading control GAPDH and expressed as arbitrary units (AU). **(C)** Immunoblot analysis of CASP1 with loading control GAPDH in BMDMs primed with Pam3CSK4 for 6 hours followed by ATP treatment for 45 minutes (n = 5). **(D)** Densitometric quantification of CASP1 p20 bands in panel C. Values for CASP1 p20 are normalized to the loading control GAPDH and expressed as arbitrary units (AU). **(E)** Immunoblot analysis of CASP1 with loading control GAPDH in BMDMs primed with poly(I:C) for 6 hours followed by ATP treatment for 45 minutes (n = 5). **(F)** Densitometric quantification of CASP1 p20 bands in panel E. Values for CASP1 p20 are normalized to the loading control GAPDH and expressed as arbitrary units (AU). **(G)** Immunoblot analysis of CASP1 with loading control GAPDH in BMDMs primed with R848 for 6 hours followed by ATP treatment for 45 minutes (n = 4). **(H)** Densitometric quantification of CASP1 p20 in panel G. **(I)** Immunoblot analysis of CASP1 with loading control GAPDH in BMDMs infected with *Aeromonas dhakensis* at an MOI of 20 (n = 5). **(J)** Densitometric quantification of CASP1 p20 in panel I. Values for CASP1 p20 are normalized to the loading control GAPDH and expressed as arbitrary units (AU). Western blots are representative from at least three independent experiments. Data are shown as mean ± SEM (B, D, F, H, and I). Statistical significance was evaluated using paired *t*-test. ns, non-significant (*p* > 0.05).

### TIA1 is dispensable for AIM2 and NLRC4 inflammasome activation

To test the effect of TIA1 loss on inflammasomes that do not require a priming step, we chose to measure AIM2 and NLRC4 inflammasome activations. The dsDNA ligand poly(dA:dT) is an activator of AIM2 inflammasome (28,29). Similarly, *Salmonella enterica* serovar Typhimurium has been shown to activate NLRC4 (11,30). No difference in the level of CASP1 activation in WT and *Tia1*^-/-^ BMDMs transfected with poly(dA:dT) (**Fig. 4A-B**) or infected with wild type *S.* Typhimurium (**Fig. 4C-D**) was observed suggesting TIA1 does not regulate AIM2 and NLRC4 inflammasome activation.

**Fig 4:**
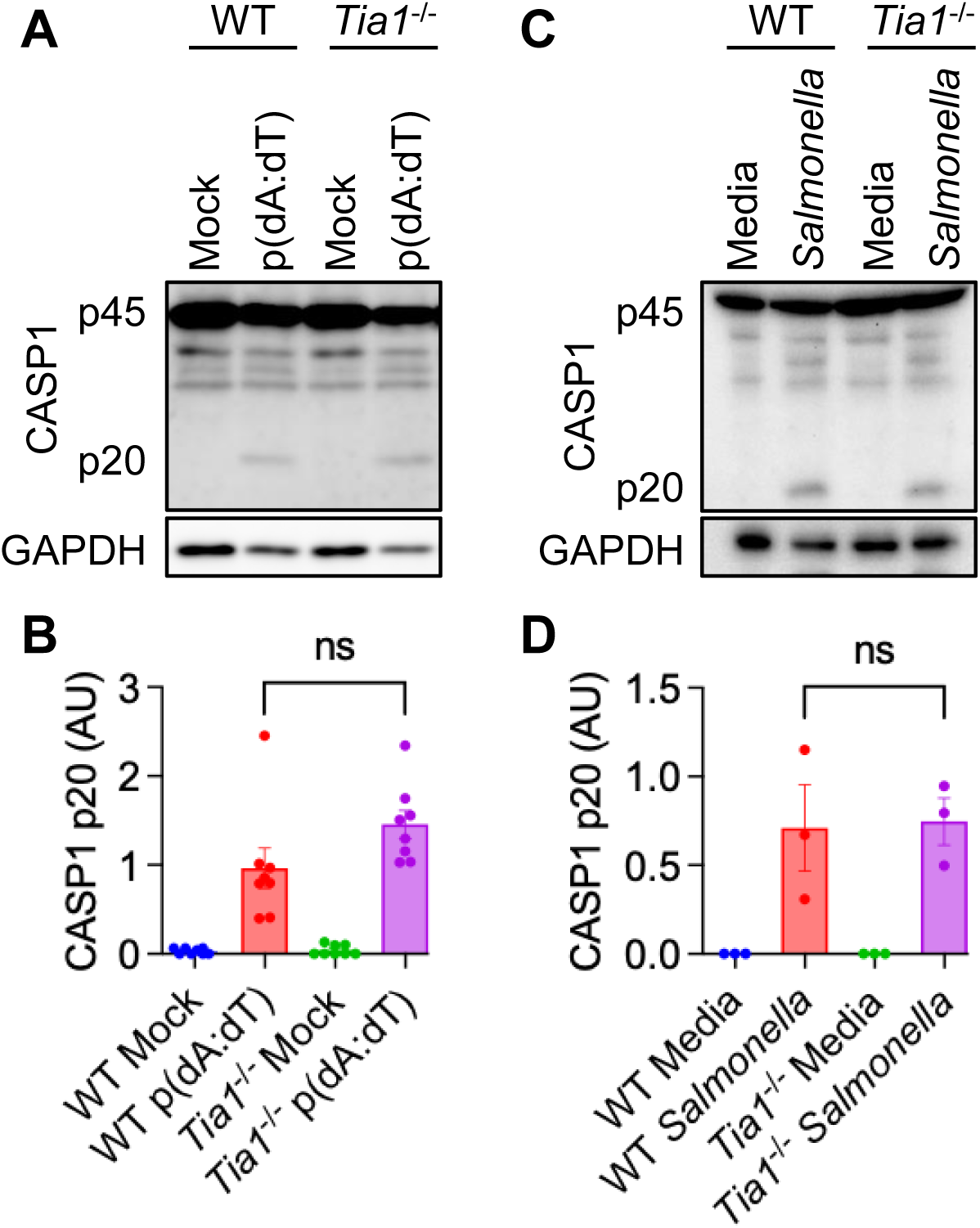
TIA1 is dispensable for AIM2 and NLRC4 inflammasome activation. **(A)** Immunoblot analysis of CASP1 with the loading control GAPDH in BMDMs transfected with poly(dA:dT) for 2 hours (n = 8). **(B)** Densitometric quantification of CASP1 p20 bands in immunoblot in panel A. Values for CASP1 p20 are normalized to the loading control GAPDH and expressed as arbitrary units (AU). **(C)** Immunoblot analysis of CASP1 with the loading control GAPDH in BMDMs infected with *Salmonella enterica* serovar Typhimurium at an MOI of 1 (n = 3). **(D)** Densitometric quantification of CASP1 p20 bands in immunoblot in panel C. Values for CASP1 p20 are normalized to the loading control GAPDH and expressed as arbitrary units (AU). Western blots are representative of at least three independent experiments. The data are shown as mean ± SEM (B, and D). Statistical significance was evaluated using paired *t*-test. ns, non-significant (*p* > 0.05).

### TIA1 does not regulate expression of core inflammasome components during non-canonical NLRP3 inflammasome activation

One way to increase non-canonical NLRP3 inflammasome activation would be to increase abundances of its critical components. Pre-treatment of macrophages with TLR agonists boosts expression of core inflammasome components (31). Therefore, we investigated how TIA1 loss impacted gene expression of core inflammasome components. First, we validated loss of TIA1 in *Tia1*^-/-^ BMDMs by reverse-transcription qPCR (RT-qPCR). Expression of *Tia1* mRNA is almost undetectable in *Tia1*^-/-^ BMDMs (**Fig. 5A, G**). Interestingly, the *Tia1* mRNA is significantly reduced in WT BMDMs transfected with LPS (**Fig. 5A, G**) suggesting that TIA1 inhibition promotes optimal inflammasome activation. Further, we evaluated the mRNA abundance of *Casp1*, *Nlrp3*, *Casp11*, *Asc* and *Il1b* genes. In response to Pam3CSK4 plus LPS transfection or LPS transfection alone, the mRNA expression of *Nlrp3* (**Fig. 5B, H**), *Asc* (**Fig. 5C, I**), *Casp1* (**Fig. 5D, J**), *Casp11* (**Fig. 5E, K**) and *Il1b* (**Fig. 5F, L**) genes were comparable between WT and *Tia1*^-/-^ BMDMs. As expected Pam3CSK4 priming boosted expression of all these genes except for *Asc*. Overall, our data suggest that TIA1 does not regulate the expression of inflammasome components during non-canonical inflammasome activation.

**Fig 5:**
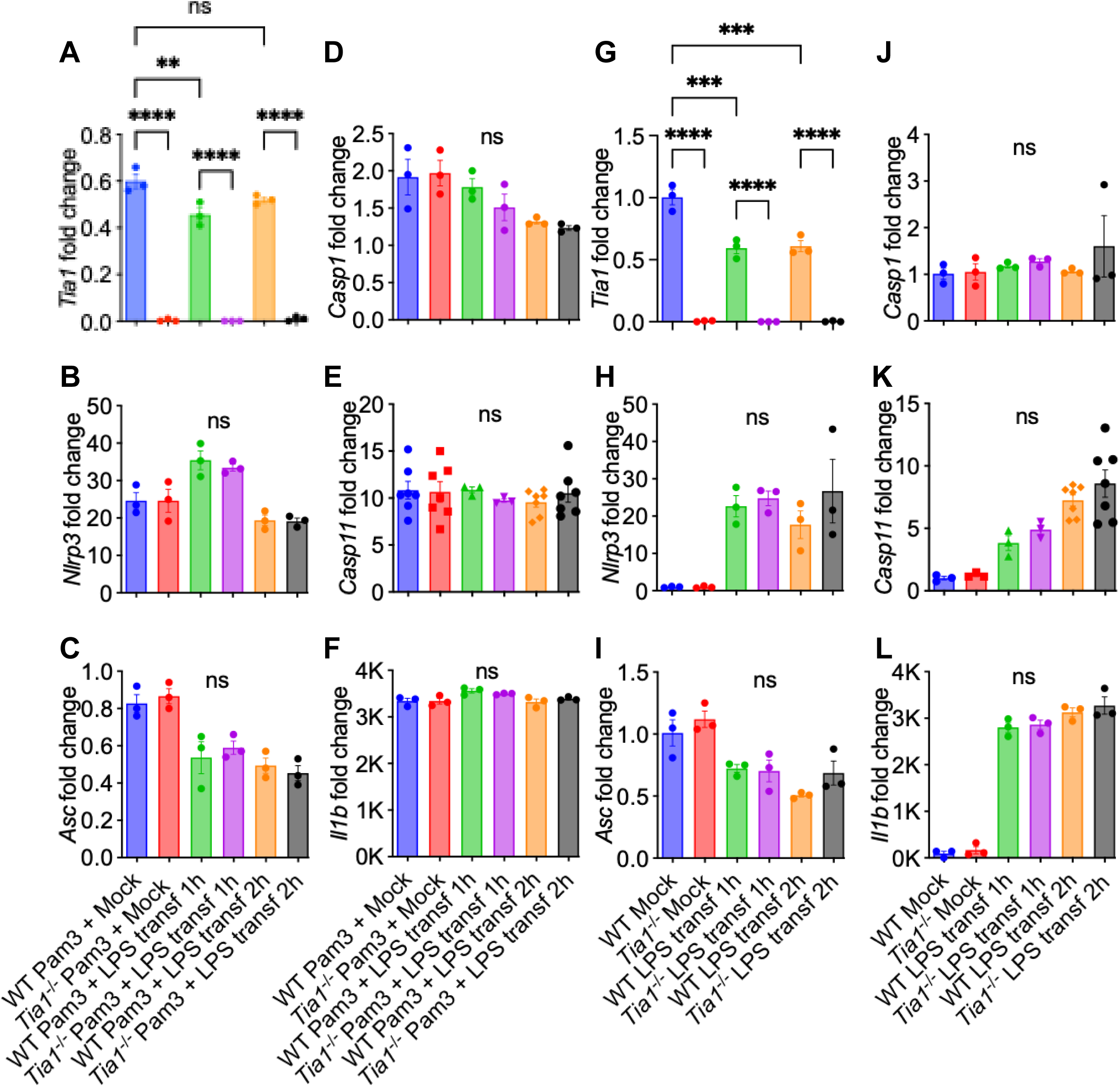
TIA1 does not regulate gene expression of core inflammasome components during non-canonical NLRP3 inflammasome activation. **(A-F)** Expression of *Tia1* (A), *Nlrp3* (B), *Asc* (C), *Casp1* (D), *Casp11* (E), and *Il1b* (F) in BMDMs primed with Pam3CSK4 for 2 hours followed by LPS transfection for 1 hour and 2 hours. (**G-L)** Expression of *Tia1* (G), *Nlrp3* (H), *Asc* (I), *Casp1* (J), *Casp11* (K), and *Il1b* (L) in BMDMs following LPS transfection for 1 hour and 2 hours. Data are from at least three independent experiments. The data are shown as mean ± SEM (A-L). Fold changes are calculated against expression in the mock treated condition. Statistical significance was evaluated using two-way ANOVA with Tukey’s multiple comparison test. ns, non-significant (*p* > 0.05), ∗∗*p* < 0.01, ∗∗∗*p* < 0.001, and ∗∗∗∗*p* < 0.0001.

### TIA1 does not regulate gene expression of core inflammasome components during canonical NLRP3 inflammasome activation

To evaluate the effect of TIA1 loss on gene expression of inflammasome components, we analyzed mRNA abundances in WT and *Tia1*^-/-^ BMDMs after priming with the canonical NLRP3 inflammasome using LPS, Pam3CSK4, poly(I:C) or R848. *Tia1* mRNA is significantly reduced in WT BMDMs primed with LPS (**Fig. 6A**), Pam3CSK4 (**Fig. 6G**), poly(I:C) (**Fig. 6M**), and R848 (**Fig. 6S**) confirming *Tia1* expression is repressed during inflammasome priming. We observed similar expression of *Nlrp3* (**Fig. 6B, H, N and T**), *Asc* (**Fig. 6C, I, O and U**), *Casp1* (**Fig. 6D, J, P and V**), *Casp11* (**Fig. 6E, K, Q and W**), and *Il1b* (**Fig. 6F, L, R and X**) in WT and *Tia1*^-/-^ BMDMs after priming with LPS, Pam3CSK4, poly(I:C) or R848. Taken together, our data suggest that loss of TIA1 has no impact on expression of genes coding for core components of the canonical NLRP3 inflammasome.

**Fig 6:**
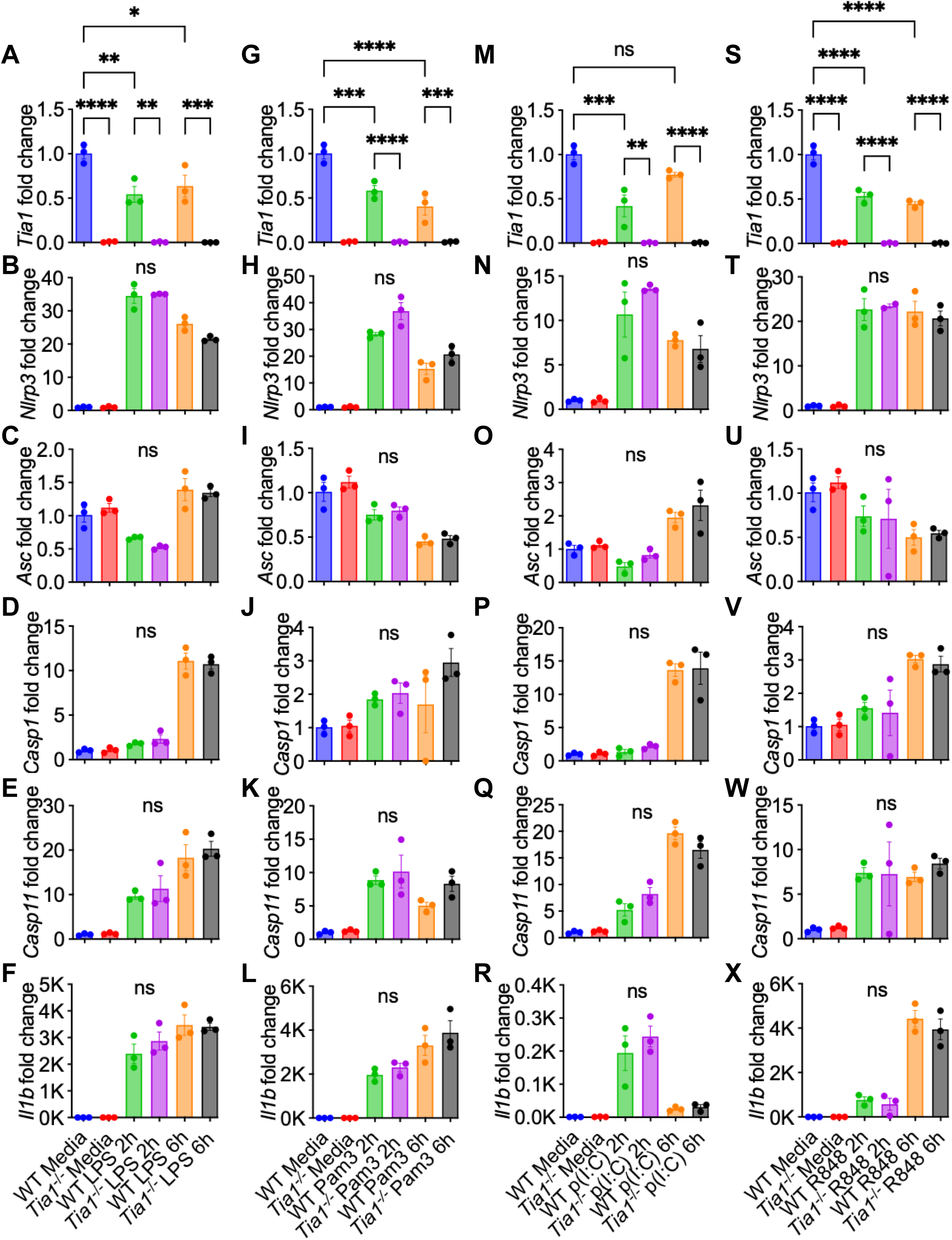
TIA1 does not regulate gene expression of core inflammasome components during canonical NLRP3 inflammasome activation. **(A-F)** Expression of *Tia1* (A), *Nlrp3* (B), *Asc* (C), *Casp1* (D), *Casp11* (E), and *Il1b* (F) in BMDMs primed with LPS for 2 hours and 6 hours. **(G-L)** Expression of *Tia1* (G), *Nlrp3* (H), *Asc* (I), *Casp1* (J), *Casp11* (K), and *Il1b* (L) in BMDMs primed with Pam3CSK4 for 2 hours and 6 hours. **(M-R)** The expression of *Tia1* (M), *Nlrp3* (N), *Asc* (O), *Casp1* (P), *Casp11* (Q), and *Il1b* (R) in BMDMs primed with poly(I:C) for 2 hours and 6 hours. **(S-X)** Expression of *Tia1* (S), *Nlrp3* (T), *Asc* (U), *Casp1* (V), *Casp11* (W), and *Il1b* (X) in BMDMs primed with R848 for 2 hours and 6 hours. Data are from at least three independent experiments. Data are shown as mean ± SEM (A-X). Statistical significance was evaluated using two-way ANOVA with Tukey’s multiple comparison test. ns, non-significant (*p* > 0.05), ∗*p* < 0.01, ∗∗*p* < 0.01, ∗∗∗*p* < 0.001, and ∗∗∗∗*p* < 0.0001.

### TIA1 does not regulate gene expression of Interferon (IFN), IFN inducible GTPases (GBPs), tumor necrosis factor (TNF) and a molecule (NINJ1) that induces plasma membrane rupture

Type I interferon signaling promotes non-canonical NLRP3 inflammasome activation in response to bacterial infections (12,30,32–34). Therefore, we investigated how TIA1 loss impacted expression of type I IFN and IFN inducible GBPs. In response to Pam3CSK4 plus LPS transfection or LPS transfection alone, expression of *Ifnb* (**Fig. 7A, F**), *Gbp2* (**Fig. 7B, G**), and *Gbp5* (**Fig. 7C, H**) genes were comparable between WT and *Tia1*^-/-^ BMDMs. Recently, NINJ1, a transmembrane protein, has been shown to be essential for pyroptosis related plasma membrane rupture. We found comparable abundances of *Ninj1* mRNAs (**Fig. 7D, I**) in Pam3CSK4 primed or unprimed WT and *Tia1*^-/-^ BMDMs transfected with LPS. As expected, we observed comparable expression of *Tnf* between WT and *Tia1*^-/-^ BMDM (**Fig. 7E, J**) (17). Overall, these data suggest that TIA1 does not regulate gene expression of components of the type I interferon pathway or those involved in plasma membrane rupture.

**Fig 7:**
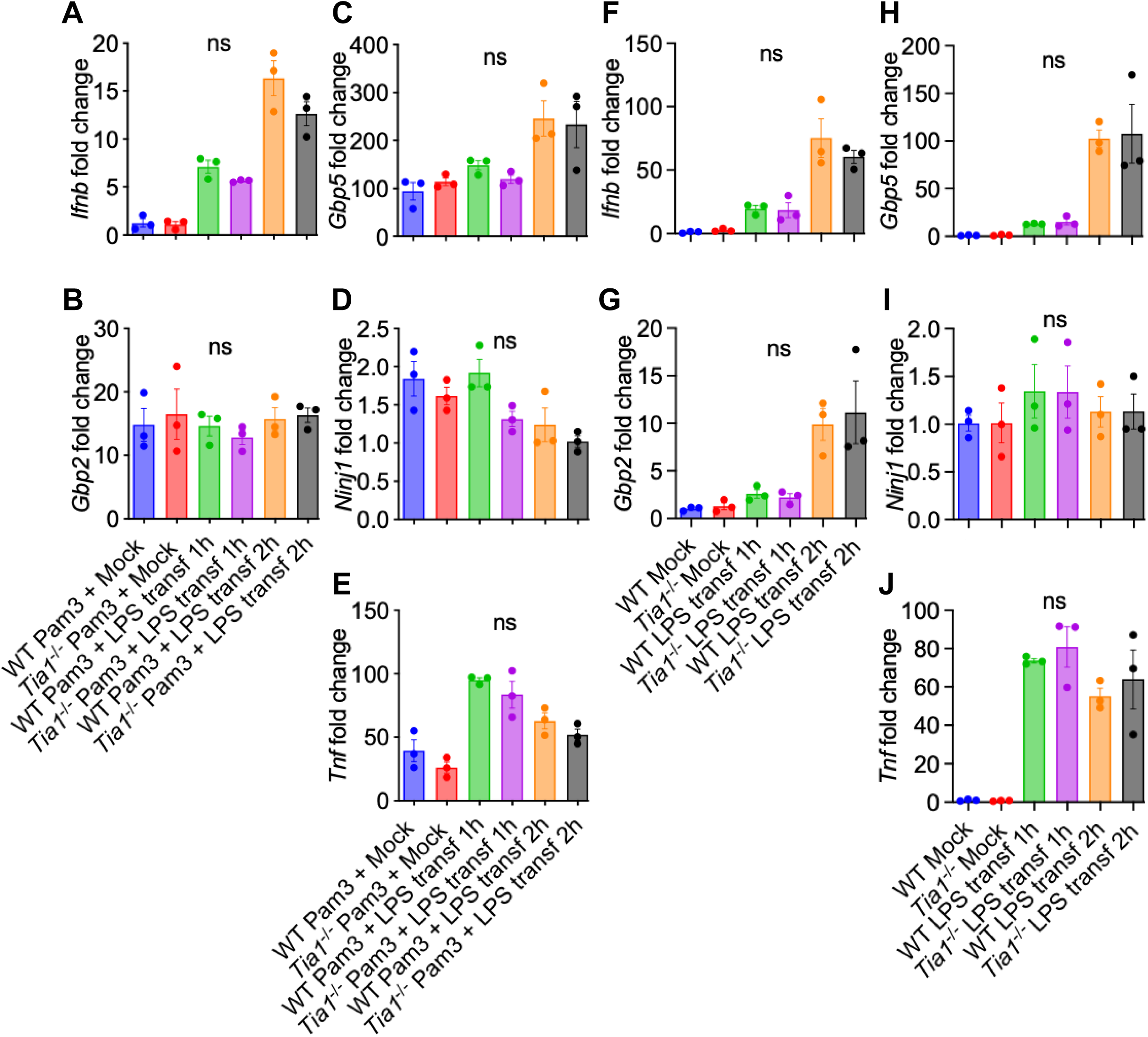
TIA1 does not regulate gene expression of type I interferon (IFN), IFN inducible GBPs, NINJ1 and TNF during non-canonical NLRP3 inflammasome activation. **(A-E)** Expression of *Ifnb* (**A)**, *Gbp2* (B), *Gbp5* (C), *Ninj1* (D), and *Tnf* (E) in BMDMs primed with Pam3CSK4 for 2 hours followed by LPS transfection for 1 hour and 2 hours. **(F-J)** Expression of *Ifnb* (F), *Gbp2* (G), *Gbp5* (H), *Ninj1* (I), and *Tnf* (J) in BMDMs following LPS transfection for 1 hour and 2 hours. Data are from three independent experiments. The data are shown as mean ± SEM (A-J). Fold changes are calculated against expression in mock treated condition. Statistical significance was evaluated using two-way ANOVA with Tukey’s multiple comparison test. ns, non-significant (*p* > 0.05).

### TIA1 does not regulate stress granules formation by non-canonical inflammasome triggers

Stress granule (SG) formation has been shown to inhibit canonical NLRP3 inflammasome activation (35). TIA1 being a key component and a nucleator of SGs (36), we investigated the possibility that SG assembly or disassembly kinetics might be responsible for inhibition of non-canonical NLRP3 inflammasome activation. We performed a time-course experiment for SG assembly and disassembly in BMDMs using G3BP1 as a SG marker with immunofluorescence confocal microscopy. We observed only a few cells with SGs after LPS transfection, with no difference between this and mock transfected BMDMs (**Fig. 8**), indicating that LPS transfection does not induce SGs. Additionally, the percentage of SG containing cells were comparable between WT and *Tia1*^-/-^ BMDMs (**Fig. 8G-H**). This indicates that TIA1 mediated inhibition of non-canonical inflammasome is independent of SG formation.

**Fig 8:**
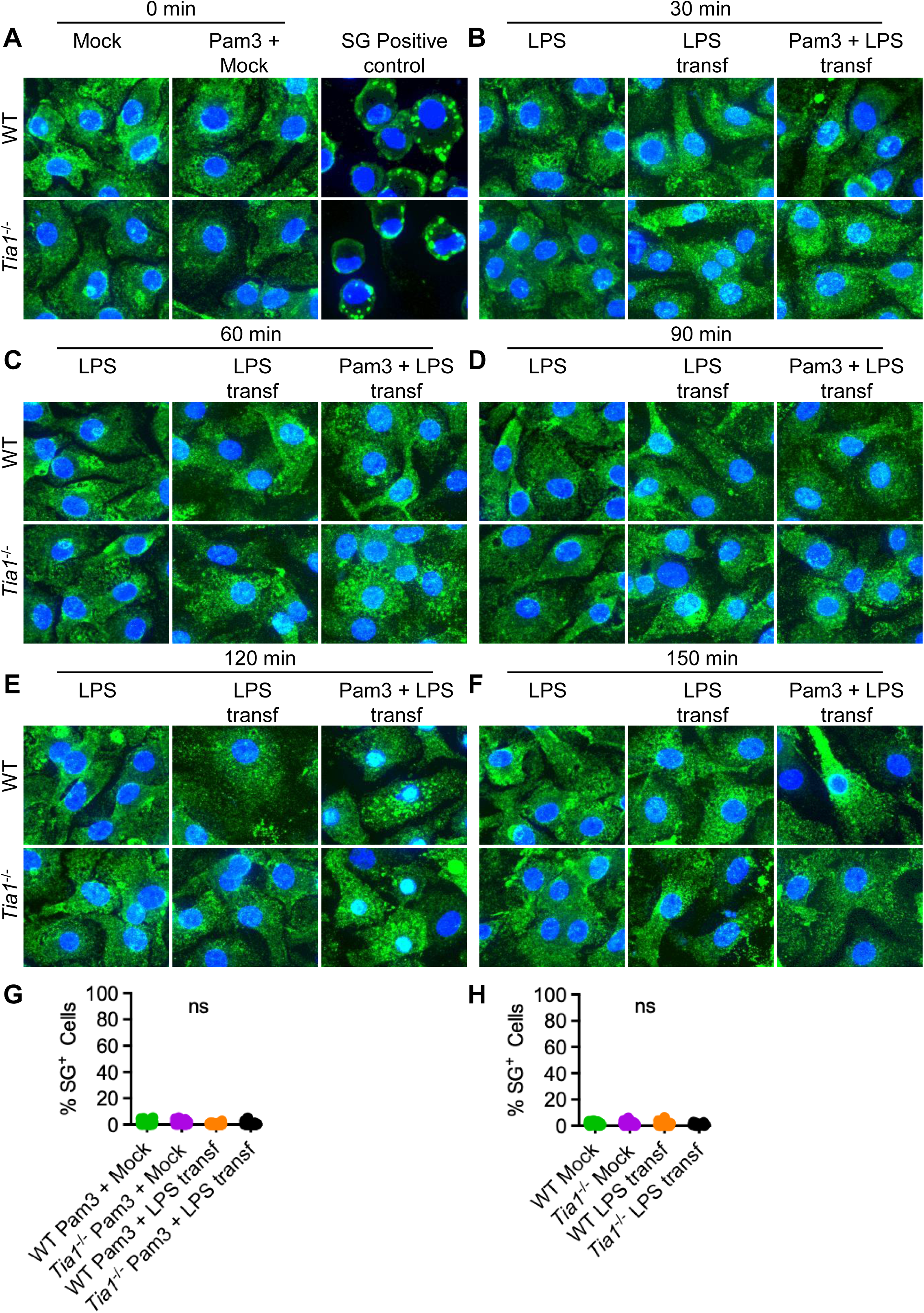
Stress granules are not assembled during non-canonical NLRP3 inflammasome activation. **(A-F)** Confocal microscopy of BMDMs primed with Pam3CSK4 for 2 hours or left unprimed (**A)** followed by treatment with LPS or transfected with LPS for 30 min **(B)**, 60 min **(C)**, 90 min **(D)**, 120 min **(E)**, and 150 min **(F)**. **(G-H)** Quantification of stress granules assembly at 150 min post LPS transfection in BMDMs primed with Pam3CSK4 (G) or left unprimed (H). Blue: DAPI, Green: G3BP1. The data are representative of at least three independent experiments. Statistical significance was evaluated using unpaired *t*-test. ns, non-significant.

### TIA1 does not regulate protein abundances of components of the canonical NLRP3 inflammasome

To test if TIA1 regulates expression of the canonical NLRP3 inflammasome components by a post-transcriptional mechanism, we analyzed protein expression by western blot. Consistent with the mRNA levels, the protein abundances of NLRP3, ASC and GSDMD were comparable between WT and *Tia1*^-/-^ BMDMs (**Fig. 9A-H**). Interestingly, the abundance of pro-CASP11 remained comparable between WT and *Tia1*^-/-^ BMDMs at basal level (**Fig. 9I-J**) or even after LPS transfection (**Fig. 9I-J**). Interestingly, Pam3CSK4 priming increased abundances of pro-CASP11 in *Tia1*^-/-^ BMDMs compared to WT BMDMs (**Fig. 9K-L**) suggesting that TIA1 might regulate TLR2 mediated post-transcriptional upregulation of pro-CASP11. However, since there was no increase in cell death (**Fig. 1M-N**) and GSDMD cleavage (**Fig. 9A, E**), increased abundance of CASP11 cannot explain observed increase in CASP1 activation and inflammasome dependent cytokine processing.

**Fig 9:**
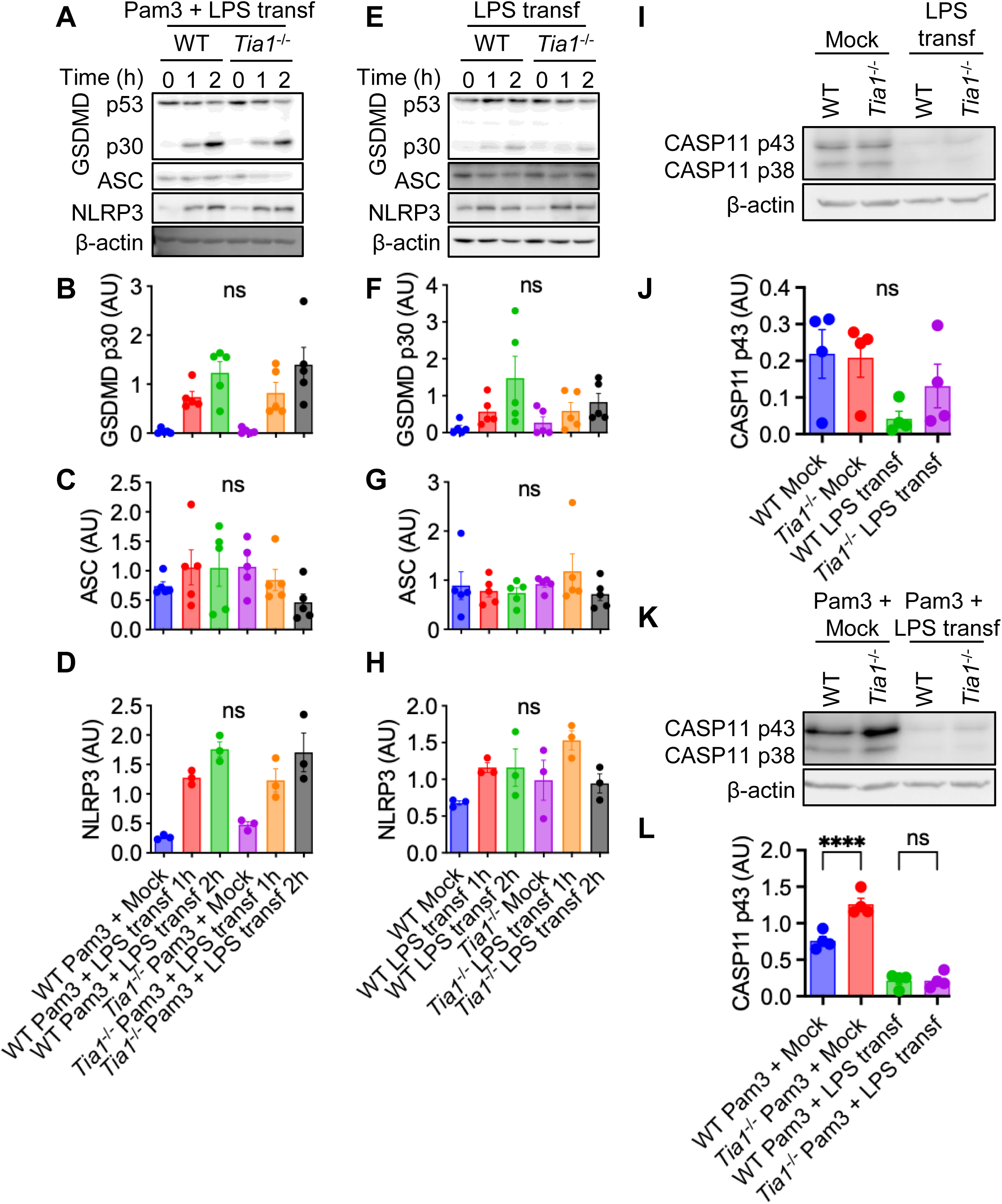
TIA1 does not regulate components of canonical NLRP3 inflammasome. **(A)** Immunoblot analysis of Gasdermin D (p50; GSDMD) and cleaved GSDMD (p30; GSDMD) (n = 5), ASC (n = 5), NLRP3 (n = 3) and loading control b-actin in BMDMs primed with Pam3CSK4 for 2 hours followed by LPS transfection for 1 hour and 2 hours. **(B-D)** Densitometric quantification of GSDMD p30, ASC, and NLRP3 bands in panel A. Values for GSDMD p30, ASC and NLRP3 are normalized to loading control and expressed as arbitrary units (AU). **(E)** Immunoblot analysis of Gasdermin D (p50; GSDMD) and cleaved GSDMD (p30; GSDMD) (n = 5), ASC (n = 5), NLRP3 (n = 3) with the loading control b-actin in BMDMs transfected with LPS for 1 hour and 2 hours. **(F-H)** Densitometric quantification of GSDMD p30, ASC, and NLRP3 bands in panel E. Values for GSDMD p30, ASC and NLRP3 are normalized to the loading control and expressed as arbitrary units (AU). **(I)** Immunoblot analysis of pro-Caspase 11 (p43; CASP11) with the loading control b-actin in Pam3CSK4 primed BMDMs transfected with LPS for 2 hours (n = 4). **(J)** Densitometric quantification of pro-CASP11 p43 bands in in panel I. Values for CASP11 p43 bands are normalized to the loading control and expressed as arbitrary units (AU). **(K)** Immunoblot analysis of pro-Caspase 11 (p43; CASP11) with the loading control b-actin in unprimed BMDMs transfected with LPS for 2 hours (n = 4). **(L)** Densitometric quantification of pro-CASP11 p43 bands in in panel K. Values for CASP11 p43 bands are normalized to the loading control and expressed as arbitrary units (AU). Western blots are representative of at least three independent experiments. The data are shown as mean ± SEM (B, C, D, F, G, H, J, and L). Statistical significance was evaluated using paired *t*-test. ns, non-significant (*p* > 0.05).

## Discussion

Septicemia is a major health care burden and is one of the leading causes of death in the United States (37,38). CASP11 and TIA1 have opposing effects on LPS mediated endotoxic shock. Animal studies suggest that CASP11 is a major molecular driver of inflammatory signaling leading to fatal septic shock (3–5,7). In contrast, TIA1 has been shown to confer protection from LPS toxicity (17). These contrasting yet related findings suggested a regulatory link between non-canonical inflammasomes and TIA1. In this study, we discovered that in response to intracellular LPS, TIA1 inhibits the non-canonical NLRP3 inflammasome which is independent of expression of major components of the complex as well as SG formation. Although these findings provide an alternative mechanistic explanation for previous observations wherein *Tia1*^-/-^ mice are hypersensitive to LPS mediated endotoxic shock, the mechanism behind our observation remains unclear (17). Interestingly, our data also showed that priming the NLRP3 inflammasome with TLR ligands is associated with a decrease in *Tia1* expression indicating that a decrease in TIA1 abundance might be important for TLR mediated inflammatory signaling. Functional significance of our observation of a decrease in *Tia1* expression during NLRP3 inflammasome priming by TLR ligands remains to be discovered.

Type I IFN signaling and IFN inducible GBPs promote non-canonical NLRP3 inflammasome activation (39). In addition, NINJ1 mediates plasma membrane rupture during pyroptosis mediated by intracellular LPS (40). Therefore, we assessed if lack of TIA1 led to increased expression of *Ifnb*, *Gbp2*, *Gbp5*, and *Ninji* in *Tia1*^-/-^ BMDMs. We did not observe an increase in expression of these genes. These findings argue against the possibility that increased non-canonical NLRP3 inflammasome activation was due to stronger type I interferon signaling.

Formation of SGs has been shown to inhibit NLRP3 inflammasome (35). TIA1 is a major nucleator of stress granules and TIA1 deficiency is linked to impaired ability to form SG (36). We did not observe SG assembly in response to LPS transfection suggesting modulation of SG is not the mechanism by which TIA1 inhibits the non-canonical NLRP3 inflammasome. Interestingly, subcellular localization of G3BP1, which is a widely used SG marker was more nuclear in Pam3CSK4 primed BMDMs after LPS transfection compared to unprimed condition. The significance of this observation requires further investigation.

There are several potential mechanisms behind TIA1 mediated inhibition of the non-canonical NLRP3 inflammasome. TIA1 might be interfering with LPS binding to CASP11. Another possibility is that TIA1 might suppress oligomerization of CASP11 induced by LPS binding through its prion-like domains (36). Post-translational modifications can control inflammasome activation (41–43). TIA1 might be regulating expression of an enzyme, for example a kinase or ubiquitin ligase, which regulates inflammasome activation. TIA1 is capable of nucleocytoplasmic shuttling. It might be inhibiting nucleocytoplasmic translocation of the inflammasome adaptor ASC. Since ASC nuclear export is required for inflammasome activation, this can explain our observed findings. Additionally, there is an increase in ASC speck positive cells in the absence of TIA1, an intriguing possibility is that TIA1 suppresses ASC speck assembly resulting in an inefficient CASP1 activation. However, if this is the case, it is not clear why TIA1 loss only affects the non-canonical NLRP3 inflammasome and this specificity needs more in-depth investigation. These and other potential mechanisms will need to be carefully evaluated in future studies.

In conclusion, we have identified TIA1 as a negative regulator of non-canonical NLRP3 inflammasome activation. Our finding extends the role of TIA1 beyond its well defined classical role as a gene expression regulator. Future studies will focus on unraveling the molecular mechanism(s) behind TIA1 mediated suppression of the non-canonical NLRP3 inflammasome while aiming towards developing drugs for treating septicemia.

## Author Contributions

PS, A and PPL designed the experiments. PPL, A and PS performed the experiments. PS supervised the study. PBK, BHN, AKC and AGT contribute with bacterial strains and experimental design. PPL wrote the first draft of the manuscript. All authors have read, edited, and agreed to the published version of the manuscript.

## Acknowledgements

We would like to thank Dr. David Place, Dr. Min Zheng, and Dr. Jian Sha for helpful discussions and suggestions about experiment design. This work was partially funded by the UTMB startup package to PS. PBK is supported by the Antimicrobial Resistance Training Program at the Texas Medical Center (T32-AI179595); Blake Neil is supported by the J.W. McLauglin predoctoral fellowship, UTMB. This research was also supported in part by the John S. Dunn Endowment funds, Institute for Human Infections, and Immunity pilot grants, and AI135453 and AI132682 NIH grants awarded to AKC. UTMB Institutional funds were awarded to AGT.

## Data Availability

All data included in the manuscript will be made available upon reasonable request to the corresponding author.

